# Insights into the evolution of oxygenic photosynthesis from a phylogenetically novel, low-light cyanobacterium

**DOI:** 10.1101/334458

**Authors:** Christen L. Grettenberger, Dawn Y. Sumner, Kate Wall, C. Titus Brown, Jonathan Eisen, Tyler J. Mackey, Ian Hawes, Anne D. Jungblut

## Abstract

Atmospheric oxygen level rose dramatically around 2.4 billion years ago due to oxygenic photosynthesis by the Cyanobacteria. The oxidation of surface environments permanently changed the future of life on Earth, yet the evolutionary processes leading to oxygen production are poorly constrained. Partial records of these evolutionary steps are preserved in the genomes of organisms phylogenetically placed between non-photosynthetic Melainabacteria, crown-group Cyanobacteria, and *Gloeobacter*, representing the earliest-branching Cyanobacteria capable of oxygenic photosynthesis. Here, we describe nearly complete, metagenome assembled genomes of an uncultured organism phylogenetically placed between the Melainabacteria and crown-group Cyanobacteria, for which we propose the name Candidatus *Aurora vandensis {au.rora* Latin noun *dawn* and *vand.ensis*, originating from Vanda}.

The metagenome assembled genome of *A. vandensis* contains homologs of most genes necessary for oxygenic photosynthesis including key reaction center proteins. Many extrinsic proteins associated with the photosystems in other species are, however, missing or poorly conserved. The assembled genome also lacks homologs of genes associated with the pigments phycocyanoerethrin, phycoeretherin and several structural parts of the phycobilisome. Based on the content of the genome, we propose an evolutionary model for increasing efficiency of oxygenic photosynthesis through the evolution of extrinsic proteins to stabilize photosystem II and I reaction centers and improve photon capture. This model suggests that the evolution of oxygenic photosynthesis may have significantly preceded oxidation of Earth’s atmosphere due to low net oxygen production by early Cyanobacteria.

## 1. Introduction

Around 2.4 billion years ago, Earth’s surface environments changed dramatically. Atmospheric oxygen rose from <10^−5^ times present atmospheric level (PAL) to >1% PAL [1-4]. This Great Oxygenation Event (GOE) permanently changed Earth’s surface geochemistry, fundamentally reshaped the cycling of key elements [5] and altered the evolutionary path of life by allowing widespread oxygen respiration [6]. The GOE was enabled by the evolution of oxygenic photosynthesis in the Cyanobacteria, making this one of the most important innovations in Earth’s history [4,7,8]. However, the evolutionary processes leading to oxygenic photosynthesis are poorly constrained [9-14]. In one hypothesis, Cyanobacteria acquired photosynthetic genes for both photosystems I and II (PSI and PSII, respectively) via horizontal gene transfer and then combined and refined them to form the photosystems that drive oxygenic photosynthesis in crown-group Cyanobacteria [15,16]. In another hypothesis, the common ancestor of all phototrophic bacteria contained the genes necessary for photosynthesis, which diversified through time and were selectively lost in non-phototrophic portions of those lineages [17-21]. In either scenario, early branching Cyanobacteria will be important to elucidating the evolution of oxygenic photosynthesis.

Due to the importance of oxygenic photosynthesis, many have attempted to extract evolutionary information by studying the genus *Gloeobacter*, the earliest branching Cyanobacteria capable of this process [22,23]1. *Gloeobacter* lack traits common in photosynthetic, non-*Gloeobacter* (crown-group Cyanobacteria) indicating that they may lack traits derived within the crown-group Cyanobacteria. For example, the *Gloeobacter* do not contain thylakoid membranes, which host photosynthesis enzymes in crown-group Cyanobacteria [24,25]. In *Gloeobacter*, photosynthesis and respiration occur in the cytoplasmic membrane [26]. *Gloeobacter* also contain a uniquely structured phycobilisome, the protein complex responsible for absorbing photons and transferring energy to the PSII reaction center. The six rods of the *Gloeobacter* phycobilisome form a single bundle whereas they are hemidiscoidal in the other crown-group Cyanobacteria [27]. Additionally, *Gloeobacter* lack PSII proteins including PsbY, PsbZ and Psb27, whereas others, including PsbO, PsbU, and PsbV, are poorly conserved [28]. As a result, *Gloeobacter* only grows slowly (23) and in low irradiance environments [29,30]. The absence of the thylakoid membrane, differences in light harvesting, and missing photosynthesis proteins help contextualize the evolution of oxygenic photosynthesis and the ecology and photochemistry of ancestral Cyanobacteria.

The Melainabacteria are an early branching sister group to the *Gloeobacter* and crown-group Cyanobacteria [10,11,31], and researchers have also interrogated their genomes for insight into the evolution of oxygenic photosynthesis [10-12,31]. Unlike the *Gloeobacter*, no known Melainabacteria have the potential for photosynthesis [10,11,31]. Therefore, the genes necessary for photosynthesis were either present in the common ancestor of Melainabacteria and Cyanobacteria and then lost in Melainabacteria and related lineages [32] or oxygenic photosynthesis evolved after the divergence of Melainabacteria and crown-group Cyanobacteria [10-12,31]. The phylogenetic space between Melainabacteria and crown-group Cyanobacteria contains an undescribed group of organisms known only from 16S rRNA gene surveys [33-36] which are either a sister group or basal to the *Gloeobacter*.

We recovered two nearly complete metagenome assembled genomes (MAGs) of a taxon within this early-diverging group from microbial mats in Lake Vanda, McMurdo Dry Valleys, Antarctica. Here, we report on the MAGs of this organism, which we have named Candidatus *Aurora vandensis*. Based on reduced photosynthetic complex within the MAG, we propose a model that sheds light on evolutionary processes that led to increased photosynthetic efficiency through stabilization of the reaction centers and better photon harvesting systems.

## 2. Methods

### Site Description

Lake Vanda is a perennially ice-covered lake located within Wright Valley, McMurdo Dry Valleys, Antarctica. Lake Vanda has a perennial ice cover of 3.5-4.0 m. The ice cover transmits 15-20% of incident photosynthetically active radiation [37]. Wavelengths shorter than 550 nm dominate the light spectrum because ice transmits little red light and water is particularly transparent to blue-green light [38]. Nutrient concentrations are low, and therefore there is little biomass in the water column [39]. However, benthic mats are abundant [38,40], covering the lake bottom from the base of the ice to >50 m [41]. The microbial mats are prostrate with abundant 0.1-30 cm tall pinnacles (41). They incorporate annual mud laminae. Mat surfaces have brown-purple coloration due to trapped sediment and pigments. The underlaying layers are characterized by green and purple pigmentation. The inner sections of large pinnacles are comprised of beige decomposing biomass. The dominant cyanobacterial genera based on morphological and 16S rRNA gene surveys are *Leptolynbya, Pseudanabaena, Wilmottia, Phormidium, Oscillatoria* and some unicellular morphotypes [42,43]. The microbial mats also contain diverse algae and other bacteria and archaea [40,44]. Incident irradiance penetrates millimeters into the mats, and most of the samples analyzed here were exposed to low irradiance in their natural environment [38].

### Sampling and DNA extraction

To obtain samples, SCUBA divers collected benthic microbial mats and brought them to the surface in sterilized plastic containers. Pinnacles were dissected in the field using sterile technique. Subsamples were placed in Zymo Xpedition buffer (Zymo Research, Irvine, CA), and cells were lysed via bead beating in the field. The stabilized samples were then frozen on dry ice and maintained frozen in the field. Samples were transported at −80 °C to UC Davis. DNA was extracted at UC Davis using the QuickDNA Fecal/Soil Microbe kit using the manufacturer’s instructions (Zymo Research, Irvine, CA, USA). The extracted DNAs were quantified using Qubit (Life Technologies) and were concentrated via evaporation until the concentration was ≥ 10 ng/uL. One bulk mat and one purple subsample were sequenced at the Joint Genome Institute (JGI).

### DNA sequencing

The JGI generated sequence data using Illumina technology. An Illumina library was constructed and sequenced 2×151 bp using the Illumina HiSeq-2500 1TB platform. BBDuk (version 37.36) was used to remove contaminants, trim reads that contained adapter sequence and right quality trim reads where quality drops to 0. BBDuk was also used to remove reads that contained 4 or more ′N′ bases, had an average quality score across the read less than 3 or had a minimum length ≤ 51 bp or 33% of the full read length. Reads mapped to masked human, cat, dog and mouse references at 93% identity were removed. Reads aligned to common microbial contaminants were also removed.

### Bioinformatic analysis

Quality controlled, filtered raw data were retrieved from IMG Gold (JGI Gold ID GP0191362 and Gp0191371). Metagenomes were individually assembled using MEGAHIT [45] using a minimum contig length of 500 bp and the paired end setting. Reads were mapped back to the assembly using Bowtie2 [46]. A depth file was created using jgi_summarize_bam_contig_depths and the assemblies were binned using MetaBAT [47]. CheckM assessed the quality of the bins [48], and bins of interest were identified based on phylogenetic placement. Average nucleotide identity (ANI) was calculated using the OrthoANI algorithm [49]. Protein coding regions were identified by prodigal [50] within CheckM. GhostKOALA and Prokka were used to annotate translated protein sequences [51,52].

When homologs of genes from the KEGG photosynthesis module were not present in the bin, they were searched for in assembled, unbinned data by performing a BLASTX search with an E-value cutoff of 1E-5. BLASTP was used to find the best hit for the retrieved sequences and to exclude those that were not the target gene. Any sequences phylogenetically similar to *A. vandensis* were identified based on their position in a phylogenetic gene tree constructed using the methodology described below.

### Phylogenetic inference

Aligned, nearly full length 16S rRNA gene sequences were collected from the Silva database (v123; [53]). The recovered 16S sequence from the bulk mat was added to this alignment using MAFFT [54]. A maximum likelihood tree was constructed in RAxML-HPC2 on XSEDE [55] in the CIPRES Science Gateway [56]. Non-full-length sequences were added to the tree using the evolutionary placement algorithm in in RAxML-HPC2 on XSEDE. Trees were rooted and visualized in the interactive tree of life [57]. Maximum likelihood trees based on 16S rRNA gene trees were separately constructed in MEGA7 [58]. For these trees, sequences were aligned with Muscle and a maximum likelihood tree was constructed using 100 bootstrap replicates.

Concatenated marker genes from Campbell et al. [59] were retrieved as described in the anvi’o workflow for phylogenomics [60]. The alignments were concatenated, and a maximum likelihood tree was constructed as described above. A maximum likelihood tree was also constructed for each individual ribosomal protein set. A genome tree was constructed in KBASE by inserting the MAGs and published Melainabacteria genomes into a species tree using the species tree builder (0.0.7; [61]).Trees were rooted and visualized in the interactive tree of life [57].

## 3. Results

Assembled metagenomes contained 313-1306 Mbp in 228837-861358 contigs with a mean sequence length of 1301-1669 bp. 49.6 and 53.3% of unassembled reads mapped back to the assembly for the bulk and purple samples, respectively. We recovered two MAGs of a taxon most closely related to *Gloeobacter,* one from each sample. The bins were 3.07 and 2.96 Mbp, had a GC content of 55.4% and 55.3%, and contained 3,025 and 3,123 protein coding sequences. Bins were 90.1 and 93.2% complete with 1.7 and 0.85% contamination based on marker gene analysis in CheckM. GhostKOALA annotated 41.1 and 41.7% of the predicted protein coding sequences. Marker gene sequences and key photosynthetic gene sequences from the bins were identical or nearly identical and the genomes were 99.96% similar based on ANI.

The MAG is most similar to *G. violaceous* with which it had 66.8% ANI across the genome. The KBASE genome tree placed the MAGs as a sister group to the *Gloeobacter* (Figure S1a). The individual marker gene trees differed in their topologies, and the concatenated tree placed *A. vandensis* as a sister group to the *Gloeobacter* (Figures 1a and S1b). The 16S rRNA gene from the MAG was >99% similar to clones from moss pillars in an Antarctic lake (AB630682) and tundra soil (JG307085) and was 91% similar to *G. violaceous* strain PCC 7421 (NR_074282; Figure 1b). Phylogenies based on 16S rRNA gene sequences varied and placed *A. vandensis* either as branching before or sister to the *Gloeobacter* dependent on which groups were included in the analysis (Figures 1 and S1c, d). The genome-based phylogeny placed *A. vandensis* as a sister group to the *Gloeobacter.*

**Figure 1:**
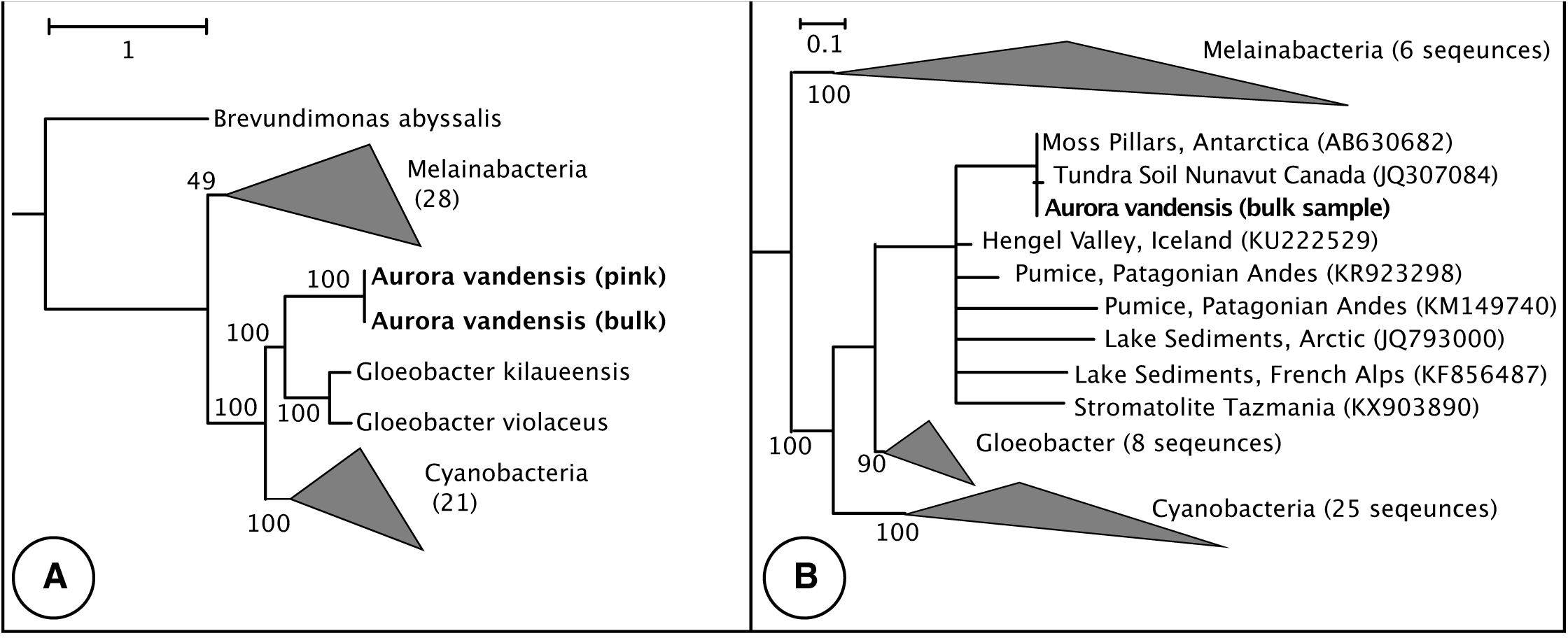
Phylogenetic placement of *A. vandensis*. A) Phylogeny constructed by inserting the *A. vandensis* bin and 38 complete or nearly complete Melainabacteria and Sericytochromatia draft genomes into a species tree containing 98 Cyanobacterial genomes. B) 16S rRNA gene phylogeny. The genus *Aurora* is indicated by the dotted line. Bootstrap values are from the original backbone tree.

Based on KEGG annotations, the MAG contained homologs of all the genes necessary for carbon fixation via the Calvin Cycle. It also contained many of the genes necessary for glycolysis via the Embden-Meyerhof-Parnas pathway (EMP; missing *pfkABC*) and citrate cycle.

The MAG contained homologs of many genes associated with oxygenic photosynthesis, but *psbJ, psbM, psbT, psbZ, psbY, psb27*, or *psbU* from photosystem II (PSII) were missing. Similarly, homologs of *psbA* were absent from the bin, but a BLASTX search of assembled, unbinned data located a *psbA* that branches before the a Gloeobacter D1 group 4 sequence and likely belongs to the MAG. PSII genes *psbP, psbO,* and *psbV* conserved (Table S1). The MAG lacked homologs of genes encoding phycobilisome proteins *apcD, apcF, cpcD, rpcG, cpcG*, and any genes associated with phycoerythrocyanin (PEC) or phycoerythrin (PE) (Table 1). The PSI genes *psaI, psaJ, psaK*, and *psaX*, and the photosynthetic electron transport gene *petJ* (cytochrome c6) were also absent. For each missing photosynthesis gene, no homologs were found in the assembled, unbinned data that had similar phylogenetic placements to other genes in the MAG, except *psbA*.

## 4. Discussion

### Genus and Species Description

We propose that our MAG is the first genome within a new genus. Compared to the most similar genome available, *G. violaceous* strain PCC 7421, it has a 66.8% average nucleotide identity (ANI) and a 91% similarity for its 16S rRNA gene. On average, genera contain taxa that are 96.5% similar based on 16S rRNA genes Therefore, we propose the creation of a new genus, *Aurora*, which includes our MAG, *Aurora vandensis,* and numerous representatives in 16S rRNA gene sequence databases. The candidate genus is named after Aurora, the goddess of the dawn, to reflect its divergence from other photosynthetic Cyanobacteria near the dawn of oxygenic photosyntheis and its presence in low light environments. Aurora also refers to the northern and southern lights aurora borealis and aurora australis, so the name also mirrors *Aurora*’s apparent preference for high latitude locations. The species, *A. vandensis,* is named after Lake Vanda where the samples originated. Lake Vanda was named after a sled dog used in the British North Greenland Expedition [62].

The phylogenetic placement of *A. vandensis* varies based on the genes or proteins used to construct the phylogeny, the taxa included in the analysis, and the tree building algorithm (e.g. Figures 1 and S1). However, it nearly always appears as sister or immediately basal to the *Gloeobacter*. Aurora’s family-level classification requires additional genomes to resolve.

To date, *Aurora* is composed of taxa from high altitude or high latitude regions including Arctic microbial mats [63], Patagonian Andes [35], Nunavut, Canada [34], The French Alps [64], and perennially ice-covered lakes in Antarctica [33] and current study; Figure 1b) and a single taxon from stromatolites in Tasmania [65]. Based on this geographic distribution, *Aurora* may be a cold adapted clade [63,66].

### Metabolic Characterization of the uncultured *Aurora* genome

*Aurora vandensis* contains homologs for the complete complement of genes necessary for carbon fixation via the Calvin Cycle and a nearly complete pathway for glycolysis via the EMP. Many Cyanobacteria contain the genes for the EMP pathway [67] and use it to ferment glycogen under dark conditions [68,69]. *Aurora vandensis* may use this pathway to ferment glycogen during the 6 months of darkness over the Antarctic winter.

*Aurora vandensis* contains homologs of many many core genes necessary for oxygenic photosynthesis, but it lacks homologs encoding several extrinsic proteins in the photosystems. As such, it is likely capable of performing oxygenic photosynthesis, but at lower efficiency than the crown-groups with more diverse extrinsic proteins.

### Photosystem II

Phycobilisomes harvest photons for use in PSII. These structures contain stacks of pigment proteins (biliproteins) connected by linker proteins and are anchored in to the thylakoid membrane in crown-group Cyanobacteria or into the cell membrane in the *Gloeobacter*. The pigments in the phycobilisome include a core of allophycocyanin (AP) which best captures photons at ∼650 nm, surrounded by rods of phycoycanin (PC; ∼620 nm), phycoerethrin (PE; maxima between 495-560 nm) and phycoerethrocyanin (PEC; 575 nm). Not all Cyanobacteria use all four pigment types, instead adapting the composition of their phycobilisomes to available irradiance [70].

*Aurora vandensis* contains homologs of the genes necessary to construct the AP core and PC rods but does not contain homologs of any biliproteins associated with PE, PEC, or many of the linker proteins associated with these pigments (Table S1). Therefore, we infer that *Aurora’s* phycobilisomes do not contain pigments that best capture energy from yellow and yellow-green photons, even though the majority of irradiance available in Lake Vanda is at wavelengths at 550 nm or below. In contrast, less than 5% of the irradiance in the AP and PC spectral ranges, which *A. vandensis* can capture, is transmitted though the ice at Lake Vanda [38].

We consider two possible hypotheses for the absence of PE and PEC related genes in *A. vandensis*: 1) presence of these genes in the common ancestor of *A. vandensis* and *Gloeobacter* and adaptive gene loss in *A. vandensis* or 2) absence in the common ancestor and addition only in the branch containing Gloeobacter and crown-group Cyanobacteria. Gene loss would limit the ability of *A. vandensis* to harvest light energy from its environment but may provide two advantages. First, because other organisms in the mat contain PE, those wavelengths are absorbed in the top few millimeters of the mat [38]. Thus, *A. vandensis* may use AP and PC to avoid competition for light with other organisms. Second, loss of PE might protect *A. vandensis* from photoinbibition. Alternately, the absence of PE in *A. vandensis* might reflect an ancestral character state of oxygenic photosynthesis with limited ability to capture photon energy. Apt *et al.,* [71] suggested that the biliproteins originated from a common ancestor, with AP being the earliest branching lineage followed by the divergence of PC and PE, and finally PEC from PC. They propose that the ancestor of all Cyanobacteria contained AP, PC, and PE biliproteins but did not contain PEC related proteins. *Aurora vandensis* partially fits this model with the absence of PEC. However, it also lacks PE. Thus, we propose an alternative model in which PE diverges after PC rather than simultaneously.

*Aurora vandensis* also lacks homologs of *apcD, acpF, cpcD*, and *rpcG*, which are structurally important to the phycobilisome and facilitate energy transfer from the antenna proteins to PSII and PSI. Knockouts of these genes in other Cyanobacteria demonstrate that they are not essential to oxygenic photosynthesis, but mutants often operate less efficiently than wildtype strains [72]. *Aurora vandensis* likely has lower effectiveness of energy transfer between the light-harvesting complex and the reaction centers relative to crown-group Cyanobacteria due to the absence of homologs of these genes. Like *Gloeobacter, A. vandensis* lacks homologs of *cpcG*, which encodes a phyobilisome rod-core linker protein. *Gloeobacter* also lacks this gene and instead uses *cpcJ* (Glr2806), which connects PC and AP, and *cpeG* (Glr1268), which connects PC and PE. These genes allow energy transfer from PC and AP to the reaction center [28,73]. *Aurora vandensis* contains sequences ∼43-58% similar to these genes, but we cannot determine if they serve the same function.

Overall, *Aurora vandensis* can likely capture irradiance for growth, but does so less efficiently than crown-group Cyanobacteria. The absence of homologs of PE creates a mismatch between available irradiance and photo capture optima, which likely limits energy transfer between the antennae proteins and the reaction centers in *A. vandensis*.

Energy flows from phocobilisomes to PSII reaction centers and excites P680, which contains the D1 and D2 reaction center dimers (*psbA* and *psbD*). This process oxidizes water and releases oxygen at the oxygen evolving complex (OEC). The reaction center also contains homologs of chlorophyll apoproteins CP43 and CP47 (*psbC* and *psbB*) and two subunits of cytochrome b559 (*psbE* and *psbF*). Other common subunits support the OEC (e.g. *psbO, psbV, psbU*) or facilitate electron flow through the reaction center.

The *A. vandensis* MAG contains homologs of all the main subunits for the PSII reaction center including the D1 and D2 proteins (Table S1). It contains homologs of *psbA* and *psbD* genes that are 91% similar to those of *G. violaceus* (WP_023172020 and WP_011142319). The translated *psbA* sequence produces a D1 protein within Group 4 [74]. Group 4 D1 proteins include all the “functional,” non-rogue D1 proteins, and all phototrophic Cyanobacteria possess a protein within this group [74].

The *A. vandensis* genome lacks a homolog of *psbM*, which helps stabilize the PSII D1/D2 dimer. However, the D1/D2 dimer still forms in the absence of PsbM in crown-group Cyanobacteria [75]. Therefore, it is unlikely that the lack of this protein prevents *A. vandensis* from forming a stable PSII reaction center. It also lacks a homolog of *psbJ*, which regulates the number of PSII reaction centers in the thylakoid membrane [76]. Mutants missing *psbJ* have less stable D1/D2 dimers and lower rates of oxygen production than wildtype strains [77]. Although *A. vandensis* may be less efficient without these genes, their absence is unlikely to prevent it from performing oxygenic photosynthesis.

When P680 reduces pheophytin a, it triggers water to donate an electron to P680 and return it to its ground redox state. Repeated four times, this process splits water into O_2_ and H^+^ at the OEC. The OEC is composed of a Mn_4_CaO_5_ cluster bound to D1, D2, CP47 and CP43 proteins. It also contains extrinsic proteins, including PsbO, PsbU, and PsbV, which help to support the OEC and provide a geochemical environment that is conducive to water oxidation [78].

The translated D1 and D2 proteins from *A. vandensis* contain all of the D1 amino acid Mn_4_CaO_5_ ligands described previously (Asp170, Glu333, Glu189, Asp342, Ala344, His332, His 337, and Ala344; [79] and the D2 Glu69 ligand [80]. The gene encoding PsbO is poorly conserved in *A. vandensis* and is only 46% similar similar to PsbO in *Gloeobacter* and 36% or less similar to those in other crown-group Cyanobacteria compared with ∼55% or greater similarity among crown-group Cyanobacteria. Despite this, PsbO in *A. vandensis* contains all the features necessary to interact with other PSII proteins and the D1, D2, CP43 and C47 subunits [81]. Therefore, the *A. vandensis* PsbO likely helps stabilize the Mn_4_CaO_5_ cluster and support the OEC despite the lack of sequence similarity. Similarly, PsbV in *A. vandensis* is dissimilar to that in crown-group Cyanobacteria, *Synechocystis* sp. PCC 6803 mutants that lack this gene are capable of evolving oxygen [82,83]. *Aurora vandensis* appears to be missing homologs of a gene encoding PsbU which stabilizes the OEC [84]. Cyanobacterial mutants missing *psbU* have decreased energy transfer between AP and PSII [85], are highly susceptible to photoinhibition, have decreased light utilization under low-light conditions, and have lowered oxygen evolution and electron donation rates than the wildtype [86]. In addition, the OEC becomes significantly more labile [86].

PsbO, PsbU, PsbV, a region of the D1, and other extrinsic proteins help control the concentration of Cl^-^, Ca^2+,^ and H^+^ and create an environment that is amenable to water oxidation [87-89]. Specifically, chloride may be involved in removing protons from the OEC [90]. Although PsbU is missing in *A. vandensis*, the other proteins conserve important residues. For example, the D1 chloride ligand site Asn338 is conserved in the translated *psbA*, but the sequence is not long enough to determine if Glu354 is also conserved. Similarly, the translated *psbO* contains Glu54, Glu114, and His231 residues that bind with Ca^2+^ [91], suggesting some Cl^-^ and Ca^2+^ regulation capabilities in *A. vandensis*.

### Cytochrome b6f

Once through PSII, the electrons move through an electron transport chain, and pass through the cytochrome b6f complex, which pumps protons across the membrane. This process creates a proton gradient that is used to generate ATP. The cytochrome b6f complex is composed of eight subunits. The *A. vandensis* genome contains homologs of genes encoding five of these subunits, including the four large subunits, PetA, PetB, PetC, PetD, PetM and the small subunit PetG. However, it appears to be missing *petL* and *petN*. A *Synechocystis* mutant was able to grow photoautotrophcally without *petL* but the rate of oxygen evolution was reduced [92]. Deletion of *petN* prevents plants from photosynthesizing [93,94]. These results have been interpreted to mean that *petN* is necessary for photosynthesis in plants and Cyanobacteria [92,95] but attempts to delete *petN* in Cyanobacteria have been unsuccessful [92] so it is not possible to determine what effect its absence may have on electron transport in *A. vandensis*. Overall, the absence of these genes may cause *A. vandensis* to transfer energy less efficiently than other Cyanobacteria but likely does not prohibit it from performing oxygenic photosynthesis or aerobic respiration.

Cytochrome b6f is restricted to crown-group Cyanobacteria, *Gloeobacter*, and *Aurora*. The Melainabacteria and Sericytochromatia contain multiple aerobic respiratory pathways, but do not contain cytochrome b6f. This has been interpreted as evidence that these three classes independently acquired aerobic respiration [12]. Based on the presence of cytochrome b6f in *Aurora* we infer that aerobic respiration evolved before the divergence of *Aurora* from *Gloeobacter,* and thus the ability to perform oxygenic photosynthesis also predated this divergence.

### Photosystem I

The end of the electron transport chain is either plastocyanin or cytochrome c6, which donate electrons to P700 in PSI. *Aurora vandensis* contains homologs of genes necessary to produce plastocyanin, but lacks homologs of *petJ,* which codes for cytochrome c6, so plastocyanain is the final electron carrier delivering electrons to PSI in *A. vandensis*.

Photosystem I in *A. vandensis* is similar to that in *Gloeobacter*. Both contain all the main subunits for PSI, but lack homologs of several genes including *psaI, psaJ, psaK*, and *psaL* that are present in crown-group Cyanobacteria. In addition, both contain homologs of many genes involved in chlorophyll biosynthesis. Therefore, PSI in *A. vandensis* likely functions similarly to PSI in *Gloeobacter*

### Photoprotection

Cyanobacteria can experience photoinhibition under high light conditions when photon absorption outstrips the ability to dissipate electrons through photochemical pathways, and reactive oxygen species accumulate at the PSII reaction center. These reactive species damage photosynthetic machinery, especially the D1 protein, which requires reassembly proteins (96-98). Cyanobacteria protect themselves from photoinhibition in two key ways. First, they use orange carotenoid proteins (OCP) as receptors to reduce the amount of energy transferred from the phycobilisome to PSII and PSI [96]. The *A. vandensis* genome contains two copies of a gene coding for a protein 68% similar to the OCP in *G. violaceous.* The OCP interacts directly with the phycobilisome [96]. Thus, the sequence differences may reflect structural differences in the phycobilisomes of *A. vandensis* and *G. violaceous.*

Cyanobacteria also protect themselves from photoinhibition using high light inducible proteins (HLIP) to dissipate energy. *Aurora vandensis* contains homologs of genes for three proteins that are 69-85% similar to HLIP in *G. violaceous.* We hypothesize that these genes act as HLIP and protect *A. vandensis* against photoinhibition.

Despite containing mechanisms for photoprotection, *A. vandensis* occupies a low-irradiance environment in Lake Vanda, particularly in the wavelengths absorbed by its biliproteins. Similarly, many other *Aurora* taxa originated from low irradiance environments. For example, one was collected from Hotoke-Ike where only 20-30% of incident PAR reaches the lake bed [97]. 16S rRNA gene sequences were found at 1 cm depth in sediments [98] where they were protected from light. Additionally, biomass may shield *Aurora* from irradiance in soil crusts in Greenland [36]. *Gloeobacter* are also sensitive to high irradiance [24] and if both *Gloeobacter* and *A. vandensis* are low-light adapted, this may be an ancestral trait of the Cyanobacteria.

### Conceptual model of the evolution of Cyanobacteria and photosynthesis

The exact phylogenetic placement of *Aurora* is uncertain and diverged before the divergence of *Gleobacter* and crown-group Cyanobacteria or is a sister group to the *Gloeobacter*. *Aurora vandensis* lacks many of the photosynthetic genes present in photosynthetic Cyanobacteria which may resemble the gene content of the ancestor of it and other Cyanobacteria. Based on these traits, we propose a model for progressive evolutionary stabilization of early oxygenic photosynthesis. Alternative models calling on gene loss or horizontal gene transfer (HGT) can also explain differences among *Aurora, Gloeobacter* and crown-group Cyanobacteria (Figure 2b, c).

**Figure 2:**
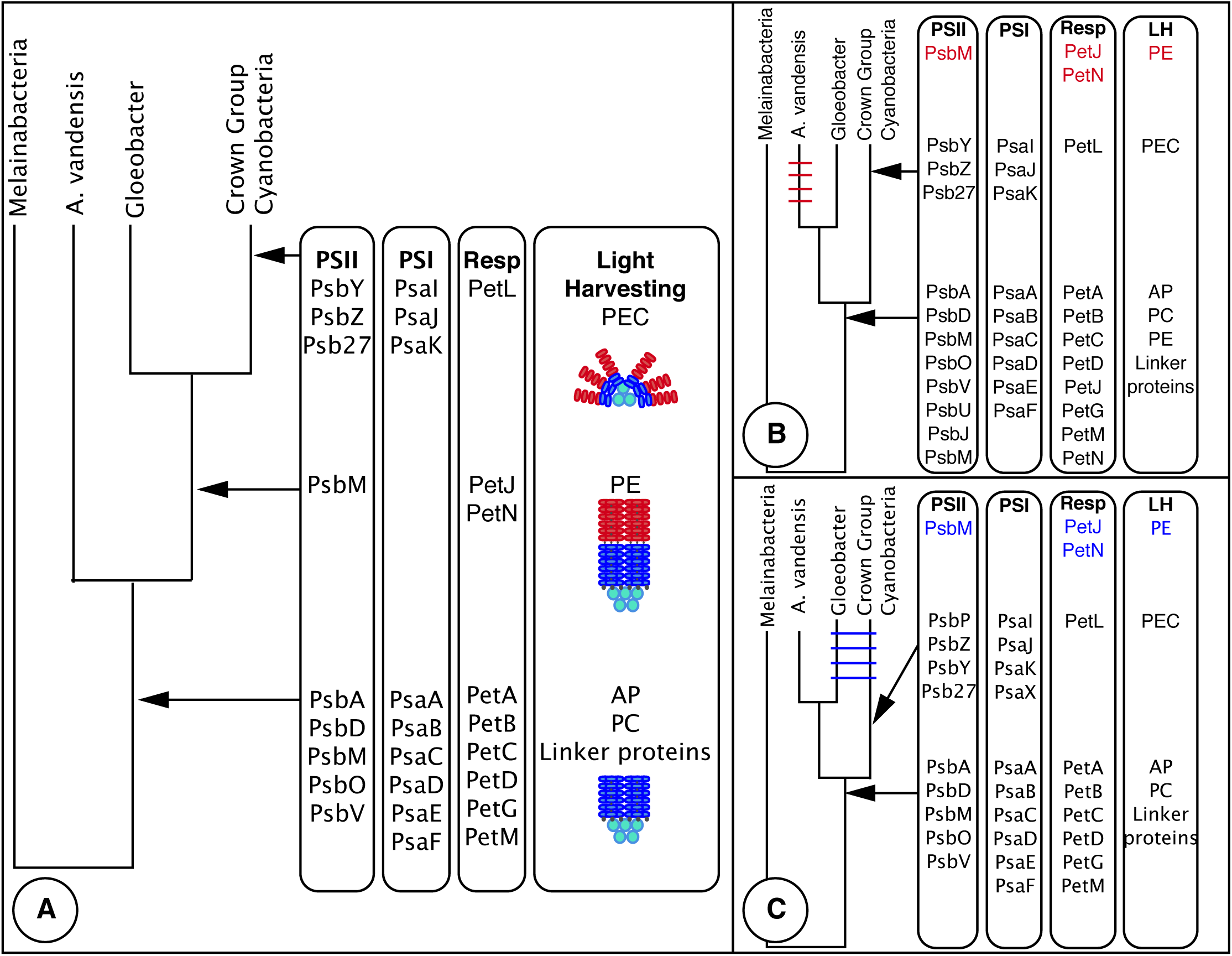
Evolutionary model of oxygenic photosynthesis. A) Our preferred model showing the progressive stabilization of oxygenic photosynthesis through time with *Aurora* basal to the *Gloeobacter.* B) Model showing gene loss in the genus *Aurora.* C) Model showing horizontal gene transfer between the ancestor of *Gloeobacter* and the ancestor of crown-group Cyanobacteria. Models B and C show *Aurora* as a sister clade to *Gloeobacter*.

For the progressive evolutionary stabilization model core photosynthetic domains were present in Cyanobacteria prior to the divergence of *Aurora* and *Gloeobacter* and were stabilized and became more efficient through the course of evolutionary time in some lineages (Figure 2a). This model predicts that the common ancestor of *Aurora, Gloeobacter*, and crown-group Cyanobacteria contained genes encoding core photosynthetic proteins including PsbA, PsbD, PsaA, PsaB, extrinsic proteins including PsbO, PsbM, and PsbV, and the AP and PC biliproteins (Figure 2a). Many of these genes appear to be essential for photosynthesis and were likely present in the common ancestor of all oxygenic phototrophs, possibly before PSII and PSI were linked to perform oxygenic photosynthesis. After the divergence of *Aurora* from *Gloeobacter* and crown-group Cyanobacteria, extrinsic proteins evolved to stabilize the reaction centers, improve water splitting, improve the flow of electrons through the reaction centers, and aid in the assembly of the reaction center. The lineage also expanded its ability to capture photons with the evolution of PE (Figure 2a). Finally, between the divergence of *Gloeobacter* and diversification of crown-group Cyanobacteria, additional extrinsic proteins were added to PSII, PEC was added to the phycobilisome, and PsaIJK and PsaX were added to PSI (Figure 2a). These reflect continued stabilization, and many may have been associated with the evolution and stabilization of the thylakoid membrane. In this model, each protein addition is predicted to increase the efficiency of oxygenic photosynthesis and be driven by selection processes.

Many alternative evolutionary models exist that rely on gene loss or HGT to explain the distribution of photosynthetic genes in *Aurora, Gloeobacter*, and crown-group Cyanobacteria. End-members models include one that relies exclusively on gene loss and another that relies on HGT (Figures 2b, c). In both models, core and extrinsic photosystems genes and much of the ETC and aerobic respiratory pathways were present in the common ancestor of the crown-group Cyanobacteria, *Aurora*, and *Gloeobacter* (Figure 2b, c). This organism also possessed AP, PC, and linker proteins for the phycobilisome. In the gene loss model, the common ancestor also contained the genes for additional extrinsic proteins in PSII, PsaIJK and PsaX in PSI, and PE. These genes were then lost in *Aurora* (Figure 2b). In the HGT model, this suite of genes evolved independently either within the *Gloeobacter* or between the divergence of *Gloeobacter* and the diversification of crown-group Cyanobacteria. The genes were then transferred between these two groups, but not into *Aurora*. Horizontal transfer appears more parsimonious than gene loss because a single HGT event can transfer multiple photosynthetic genes [99,100] and the transfer of beneficial traits between *Gloeobacter* and crown-group Cyanobacteria seems more likely than their loss.

*Aurora* branches before the divergence of *Gloeobacter* and crown-group Cyanobacteria (Figure 2a) is most parsimonious with the emergence of oxygenic photosynthesis, a new metabolism capable of generating large amounts of chemical energy from light energy but at the expense of significant metabolic machinery damage. Through time, evolutionary pressures led to progressive increases in stability and productivity in some lineages, which allowed the expansion of early Cyanobacteria into environments with greater irradiance. Based on this model, we predict that ancestral lineages that emerged prior to the GOE may have needed to occupy low irradiance habitats due to photoinhibition, and high UV doses that would have accompanied other wavelengths in the pre-oxygenated atmosphere. As the photosystems stabilized, photon capture efficiency improved, and oxygenic phototrophs expanded to higher-light environments. Both would have resulted in significantly higher primary productivity and rates of oxygen production.

### Importance of *Aurora vandensis*

The crown-group Cyanobacteria diversified between 2.3 and 1.9 billion years ago [14], approximately 600 to 900 million years after the divergence of the phototrophic Cyanobacteria and the Melainabacteria [14]. The only characterized lineages that diverged within this interval are *G. violaceous, G. kilauensis*, which diverged 2.2 to 2.6 billion years ago, and now *A. vandensis,* with *A. vandensis* potentially diverging between the Melainabacteria and *Gloeobacter*. If basal to the *Gloeobacter,* this new genome provides key insight into the evolutionary processes occurring over the 300-650 million years [14,104] spanning the invention of the most transformative metabolism on Earth, oxygenic photosynthesis. Thus, the genome of *A. vandensis* is particularly important for contextualizing this innovation. Specifically, an evolutionary model in which *Aurora* is basal to *Gloeobacter* (Figure 2a) is parsimonious with the emergence of oxygenic photosynthesis as a new metabolism capable of generating substantial chemical energy from light but at the expense of significant metabolic machinery damage. Thus, early cyanobacterial lineages may have inhabited only low irradiance habitats due to photoinhibition. Through time, evolutionary selection led to progressive increases in stability and productivity, which allowed expansion of Cyanobacteria into environments with greater irradiance. As photosystems stabilized, photon capture efficiency also improved, increasing primary productivity. Eventually, habitat expansion and improvements in efficiency allowed Cyanobacteria to produce enough oxygen to cause oxidative weathering [101] and finally trigger the GOE [3].

Low photosynthetic efficiency in early Cyanobacteria can reconcile models that predict rapid oxidation of Earth’s surface [102] with the geological record, which shows whiffs of oxygen before the GOE [4,103]. If our evolutionary model is correct, cyanobacterial oxygen production could have initiated long before oxygen accumulated in the oceans and environment.

## Acknowledgements

Sequencing was provided by the U.S. Department of Energy Joint Genome Institute, a DOE Office of Science User Facility, and is supported under Contract No. CSP502867. Samples used in this project were collected during a field season supported by the New Zealand Foundation for Research, Science and Technology (grant number CO1×0306) with field logistics provided by Antarctica New Zealand (project K-081). Salary support for CG was provided by the Massachusetts Institute of Technology node of the NASA Astrobiology Institute.

---

**Figure S1.** A) Genome phylogeny from KBASE showing *A. vandensis* as a sister group to the *Gloeobacter.* B) 16S rRNA gene phylogeny showing the genus *Aurora* as basal to the *Gloeobacter*. C) Ribosomal protein L2 phylogeny with *A. vandensis* sister to the *Gloeobacter* and D) IF3 C phylogeny showing *A. vandensis* diverging before the divergence of the *Gloeobacter*.

**Table S1.** Photosynthetic genes present in *A. vandensis, Gloeobacter*, and crown-group *Cyanobacteria*. Differences between the early branching Aurora and Gloeobacter and the crown-group Cyanobacteria are indicated in green. Difference between Aurora and Gloeobacter are indicated in blue. Modified from ref 8.

